# Early Deficits in Dentate Circuit and Behavioral Pattern Separation after Concussive Brain Injury

**DOI:** 10.1101/2023.06.22.546120

**Authors:** Lucas Corrubia, Andrew Huang, Susan Nguyen, Michael W. Shiflett, Mathew V. Jones, Laura A. Ewell, Vijayalakshmi Santhakumar

**Affiliations:** Department of Pharmacology, Physiology and Neuroscience, Rutgers New Jersey Medical School, Newark, New Jersey 07103; Department of Molecular, Cell and Systems Biology, University of California Riverside, Riverside, California 92521; Department of Psychology, Rutgers University, Newark, NJ 07102; Department of Neuroscience, University of Wisconsin, Madison, WI, 53705; Department of Anatomy and Neurobiology, University of California Irvine, Irvine, California 92697

**Keywords:** granule cell, dentate gyrus, electrophysiology, traumatic brain injury, memory, behavior, fluid percussion injury, temporal pattern separation

## Abstract

Traumatic brain injury leads to cellular and circuit changes in the dentate gyrus, a gateway to hippocampal information processing. Intrinsic granule cell firing properties and strong feedback inhibition in the dentate are proposed as critical to its ability to generate unique representation of similar inputs by a process known as pattern separation. Here we evaluate the impact of brain injury on cellular decorrelation of temporally patterned inputs in slices and behavioral discrimination of spatial locations *in vivo* one week after concussive lateral fluid percussion injury (FPI) in mice. Despite posttraumatic increases in perforant path evoked excitatory drive to granule cells and enhanced ΔFosB labeling, indicating sustained increase in excitability, the reliability of granule cell spiking was not compromised after FPI. Although granule cells continued to effectively decorrelate output spike trains recorded in response to similar temporally patterned input sets after FPI, their ability to decorrelate highly similar input patterns was reduced. In parallel, encoding of similar spatial locations in a novel object location task that involves the dentate inhibitory circuits was impaired one week after FPI. Injury induced changes in pattern separation were accompanied by loss of somatostatin expressing inhibitory neurons in the hilus. Together, these data suggest that the early posttraumatic changes in the dentate circuit undermine dentate circuit decorrelation of temporal input patterns as well as behavioral discrimination of similar spatial locations, both of which could contribute to deficits in episodic memory.

## Introduction

Traumatic brain injury (TBI) is a growing epidemic (Vaishnavi et al., 2009; Rusnak, 2013) with an annual global incidence of over 60 million patients (McAllister, 1992) who are at risk for a variety of neuropsychological sequelae (Rao and Lyketsos, 2000). While neurological complications of TBI increase with injury severity (Annegers and Coan, 2000), even mild to moderate concussive brain injuries can lead to early and long term deficits in memory function (Kumar et al., 2013; Arciniega et al., 2021; Alosco et al., 2023). The dentate gyrus, a critical node in memory processing, receives extensive inputs from the entorhinal cortex and develops discrete sparse representations for downstream processing in hippocampal circuits (Leutgeb et al., 2007; Neunuebel and Knierim, 2014). The dentate gyrus is also the focus of neuronal loss and circuit reorganization early after human and experimental concussive brain injury (Lowenstein et al., 1992; Santhakumar et al., 2000; Meier et al., 2016; Leh et al., 2017; Neuberger et al., 2017b; Folweiler et al., 2020), suggesting that dentate dependent memory functions are likely compromised after brain injury.

In particular, the dentate gyrus has been shown to be essential for distinguishing between similar memory episodes by encoding partially overlapping memories into discrete neuronal representations (Gilbert et al., 2001; McHugh et al., 2007; Yassa and Stark, 2011; Cayco-Gajic and Silver, 2019). At the circuit level, this is proposed to be achieved by pattern separation, a process of minimizing interference by segregating similar input patterns into discrete neuronal activity patterns. Whereas behaviorally relevant inputs and the resulting neuronal input patterns occur in space and time (Hainmueller and Bartos, 2020), their neuronal representations have been evaluated mostly in the spatial domain as changes in firing or activity rates in spatially discrete neuronal groups or ensembles (Leutgeb et al., 2007; Larimer and Strowbridge, 2010; GoodSmith et al., 2017). However, whether brain injury impairs the ability of the dentate circuit to faithfully represent inputs and disambiguate similar neuronal input patterns remains unknown.

Most behavioral studies have focused on discrimination of spatial and contextual patterns to evaluate mechanisms of dentate pattern separation and assess the impact of disease (McHugh et al., 2007; Morris et al., 2012; Bekinschtein et al., 2013; Kahn et al., 2019). Studies in the fluid percussion injury (FPI) model of concussive brain injury have revealed early deficits in sequential spatial navigation in a dentate-dependent radial arm maze task, indicating that concussive brain injury compromises the ability to process and recall subtle spatial differences (Correll et al., 2021). Similarly, brain injury is also known to compromise working memory function (Smith et al., 2015; Korgaonkar et al., 2020a). Still, whether episodic processing of spatial novelty by behavioral discrimination of spatial location is altered after brain injury remains to be determined.

In an approach complementary to behavioral studies, computational analyses of mechanisms for disambiguating overlapping memories have adopted spatially and temporally patterned inputs to assess “pattern separation”. In this approach, model networks are activated by inputs with various degrees of similarity in spike timing and/or activated cell ensembles and the ability of the network to generate distinct spatio-temporal firing patterns is used as a measure of pattern separation (Rolls and Kesner, 2006; Myers and Scharfman, 2011; Madar et al., 2019b; Braganza et al., 2020). These studies have shown that dentate granule cells (GCs) decorrelate input patterns both in terms of the active ensembles recruited as well as the pattern of firing in the active neurons (Myers and Scharfman, 2009, 2011; Yim et al., 2015). However, few experimental studies have evaluated the ability of the dentate circuit to decorrelate spatially and temporally patterned inputs (Larimer and Strowbridge, 2010; Madar et al., 2019a). Recently, Madar et al. (2019a) adopted an experimental paradigm in hippocampal slices to demonstrate that GCs could undertake input decorrelation at the level of individual neurons. These studies identified a role for inhibition in supporting temporal pattern separation in GCs and revealed cell-type specific and disease related changes in input decorrelation by dentate neurons (Madar et al., 2019a; Madar et al., 2021). These findings are consistent with predictions of simulations (Braganza et al., 2020) and with behavioral deficits in discrimination of novel spatial location in mice during suppression of dentate somatostatin interneuron activity in vivo (Morales et al., 2021). Since experimental concussive brain injury has been shown to result in early loss of dentate inhibitory neuron subtypes (Lowenstein et al., 1992; Toth et al., 1997; Santhakumar et al., 2001; Folweiler et al., 2020), we examined whether the resulting changes in inhibition (Folweiler et al., 2020; Gupta et al., 2020; Gupta et al., 2022) could alter dentate dependent pattern separation in the temporal and spatial domains one week after brain injury. Because the relationship between neuronal spike train timing and behavioral spatial information is currently unclear, we will denote GC pattern separation of temporally patterned inputs in slice as “decorrelation” and behavioral spatial pattern separation in vivo as “spatial discrimination”.

## MATERIALS AND METHODS

All experiments were performed in accordance with IACUC protocols approved by the University of California, Riverside, CA in keeping with the ARRIVE guidelines.

### Fluid Percussion Injury

Randomly selected 8 to 10-week-old littermate male and female C57BL6/J mice were subjected to lateral fluid percussion injury (FPI) or sham injury. Briefly, mice were placed in a stereotaxic frame under isoflurane anesthesia and administered the local anesthetic 0.25% bupivacaine (subcutaneous). A 2.7 mm hole was trephined on the left side of the skull 2 mm caudal to bregma and 1 mm lateral from the sagittal suture. A Luer-Lok syringe hub with a 3 mm inner diameter was placed over the exposed dura and bonded to the skull with cyanoacrylate adhesive. The animals were returned to their home cage following recovery. One day later, animals were anesthetized with isoflurane and attached to a fluid percussion injury device (Virginia Commonwealth University, Richmond, VA, Model 01-B) directly at the metal nozzle. A pendulum was dropped to deliver a brief (20 ms) 1.5–1.7 atm impact on the intact dura. For sham injury, the animals underwent surgical implantation of the hub and were anesthetized and attached to the FPI device, but the pendulum was not dropped (Smith et al., 2012; Gupta et al., 2022).

### Slice Preparation

Naive C57BL6/J mice and mice one week after FPI or sham injury were anesthetized with isoflurane and decapitated for slice preparation using established protocols (Korgaonkar et al., 2020b; Afrasiabi et al., 2021). Horizontal brain slices (300 µm) were prepared in ice-cold sucrose artificial CSF (sucrose-aCSF) containing the following (in mM): 85 NaCl, 75 sucrose, 24 NaHCO_3_, 25 glucose, 4 MgCl_2_, 2.5 KCl, 1.25 NaH_2_PO_4_, and 0.5 CaCl_2_ using a Leica VT1200S Vibratome (Wetzlar, Germany). Slices were incubated at 34°C for 30 min in an interface holding chamber containing an equal volume of sucrose-aCSF and recording aCSF, and subsequently were held at room temperature. The recording aCSF contained the following (in mM): 126 NaCl, 2.5 KCl, 2 CaCl_2_, 2 MgCl_2_, 1.25 NaH_2_PO_4_, 26 NaHCO_3_, and 10 D-glucose. All solutions were saturated with 95% O_2_ and 5% CO_2_ and maintained at a pH of 7.4 for 1– 6 h. Only sections on the side of injury were used in experiments.

### Physiological Recordings

Voltage clamp recordings of perforant path evoked excitatory postsynaptic currents (eEPSCs) were obtained from dentate GCs using recording electrodes (3 - 6 MΩ) containing a Cs-Methanesulfonate Internal (in mM): 140 Cs-Methane sulfonate, 5 NaCl, 10 HEPES, 0.2 EGTA, 4 Mg-ATP, 0.2 Na-GTP, 5 Qx-314 with 0.2% biocytin. GCs were held at −70 mV, close to reversal potential for GABA, to isolate EPSCs (Andreasen and Hablitz, 1994). Evoked EPSCs were recorded in response to 0.1 to 2.0 mA stimuli, applied in 100 pA steps, through a concentric stimulus electrode positioned at the perforant path on the other side of the fissure from the dentate molecular layer. Peak amplitude of eEPSC was measured from the average of 3 to 5 responses at each step using ClampFit 10. A subset of recordings were obtained in a saturating concentration of gabazine (10 µM) to block both synaptic and extrasynaptic GABA_A_ receptors.

Current clamp recordings to assess GC pattern separation were obtained using filamented borosilicate glass electrodes (3 - 6 MΩ) containing (in mM): 126 K-gluconate, 4 KCl, 10 HEPES, 4 Mg-ATP, 0.3 Na-GTP, 10 PO-creatinine with 0.2% biocytin. To assess cell health, GCs were held at −70 mV and responses to 1500 ms current steps from −200 pA to threshold positive current injection in 40 pA steps were recorded. Cells with resting membrane potential positive to −55 mV were excluded from analysis.

### Pattern Separation Paradigm and Analysis

To assess GC pattern separation, input trains of varying correlations were adopted from Madar et al. (2019a). Each input correlation set included 5 temporally distinct trains at 10 Hz with an overall mean Pearson’s correlation coefficient (R_in_) of 0.25, 0.50, 0.75, or 0.95, designated henceforth as R25, R50, R75 and R95. The individual input trains (#1-#5) within an input correlation set were each 2 seconds long and were delivered at 2 second intervals. The 5 distinct input trains were delivered in sequence and the set was repeated 10 times for a total of 50 recorded trains (Madar et al., 2019a; Madar et al., 2021). Note that the average frequency of each input train was maintained at 10 Hz while in each set of 5 trains the timing of stimuli was varied to generate trains with different correlations.

GCs were held at −70 mV in whole cell configuration and stimuli were delivered through a concentric stimulating electrode (10 µm tip diameter) placed in the outer molecular layer to activate perforant path fibers. Stimulus trains were delivered with intensity ranging from 0.1-2 mA at 0.5 Hz so as to elicit action potentials in response to approximately 50% of stimuli. Once stimulus intensity was set, it was maintained constant for all input sets recorded. Cell resting membrane potential and access resistance were assessed between input sets and recordings were terminated if access resistance increased over 20% or membrane depolarized beyond −55 mV. To assess the role of inhibition in temporal pattern separation, GCs were recorded and stimulated with input correlations of R75 in aCSF and then perfused with a sub-saturating concentration of gabazine (SR95531, 100nM), a GABA_A_ receptor antagonist as described in earlier studies (Madar et al., 2019b), and to avoid burst firing (not shown) that occurred in pilot studies using saturating gabazine (10 µM).

For quantification of GC pattern separation function, Pearson’s R correlation was calculated on output spike train rasters as previously defined in Madar et al. (2019a) using custom MATLAB code modified from: https://github.com/antoinemadar/PatSepSpikeTrains. Unless indicated otherwise, data were binned at 10 ms. For validation of pattern separation paradigm in naïve mice, pairwise correlation of each GC spike output train (R_out_) was plotted against the predetermined correlation in the corresponding pairwise input spike trains. The five distinct input spike trains and their corresponding output spike trains are compared in a pairwise manner to assess correlation. Thus, the five input patterns yield 10 distinct R_in_ values centered around the target R_in_ and correspondingly, 10 R_out_ values. These R_in_-R_out_ sets were used to compare data between GCs from sham and FPI mice.

To assess the reliability of GC input-output coupling between experimental groups, we evaluated spike train reliability (R_w_) as a measure of the cell’s ability to faithfully respond to multiple presentations of the same input train (effectively R_in_=1). R_w_ was calculated as the average correlation of output of a given GC in response to 10 presentations of the same input pattern in the R95 input set and then averaged across the 5 patterns.

### Novel Object Location Task and Analysis

To assess behavioral pattern separation, we implemented a novel object location task to test spatial discrimination (Bekinschtein et al., 2013; Morales et al., 2021) in mice one week post FPI or sham injury. The task consisted of a circular arena 50 cm in diameter and 47 cm high with laboratory tape in varying patterns in four distinct locations on the white opaque arena walls serving as spatial visual cues. The experimenter room was separated from the testing room with the test arena by a closed door. The testing room was dimly lit with overhead lights using dimmer switch on lowest setting. Mice underwent four days of habituation for 10 minutes each followed by a sample and choice phase on day five. The testing arena and objects were cleaned with 70% ethanol in between each animal throughout the entire experiment. During habituation, mice were brought to the testing room from the vivarium, handled by the experimenter, and exposed to the testing arena for 10 min each day where they were allowed to explore freely. On test day, mice were first exposed to a sample phase for 10 min, with three identical objects placed 120 degrees and approximately 16 cm apart and allowed to freely explore the objects. Mice were then returned to their home cage for 30 min before the choice phase. During the choice phase, one object was placed in its original (familiar) location, one object was removed, and the last object was shifted by 60° to a novel (unfamiliar) location. The mice were placed into the arena and allowed to explore the objects for five minutes. Videos were manually coded and analyzed by an experimenter blinded to animal injury status. Object exploration was counted as the number of times the animal’s nose was within 0.5 cm of the object, with each continuous second counted as an additional exploration. Discrimination ratio was calculated as [Exploration of Object in Unfamiliar Location – Exploration of Object in Familiar Location] / [Total Exploration]. Total exploration was also compared between groups.

### Immunohistochemistry

Immunohistochemistry for ΔFosB, somatostatin (SST), and parvalbumin (PV) was used to evaluate persistent neuronal activity (You et al., 2017) and interneuron loss. Immediately following the novel object location task, mice were perfused transcardially with ice cold PBS followed by 4% PFA. Brains were excised and incubated in 4% PFA overnight and transferred to 30% sucrose for 2-3 days or until tissue sank. Brains were then flash frozen in OCT using liquid nitrogen and stored at −80°C prior to cryosectioning. 20 µm serial sections were collected and mounted on SuperFrost slides and stored at −80°C prior to staining. Slides were stained for ΔFosB, SST, or PV following standard procedures (Neuberger et al., 2017a; Korgaonkar et al., 2020b). Briefly, slides were allowed to thaw to room temperature (RT) for 10 min, washed with 1xPBS 3x5min to wash off residual OCT. Slides were blocked in 10% normal goat serum in PBS + 0.3% Triton-X for one hour at RT, then incubated in anti- ΔFosB, SST, or PV antibody (1:500, Cell Signaling D3S8R; SST: 1:100, Millipore MA5-16987; PV: 1:1000, Swant GP72) overnight at 4°C. Slides were then washed 3x10 min with PBS and then incubated in goat anti-rabbit AF594 secondary antibody (1:1000, Invitrogen A11012) for ΔFosB, goat anti-rat AF647 antibody (1:1000, Invitrogen A-21247) for SST goat anti-guinea pig AF647 antibody (1:1000, Invitrogen, A-21450) for PV for 1.5 hours at RT, washed 3x10 min with PBS, and coverslipped using VectaShield with DAPI mounting media. Slides were imaged using Zeiss Axioscope Epifluorescence Microscope as single optical sections at 10x and 20x for analysis. Blinded analysis was performed using Cell Detection settings in QuPath 0.4.2 (https://qupath.github.io) with files coded without injury status. Briefly, for ΔFosB and SST analysis, a ROI was drawn to outline the granule cell layer (GCL) using DAPI labeling to detect ΔFosB positive cells and in the hilus to detect SST neurons. Detection parameters were kept consistent between sections, and 3 to 6 level matched sections were averaged in each mouse for analysis, depending on slice and image quality. Robust and sparse PV immunolabeled neuronal somata were counted manually on QuPath for full dentate region to include both hilar, GCL, and molecular layer PV interneurons. SST cell counts were normalized to total hilar area (in mm^2^). The hilar area averaged across sections was not different between sections from sham and FPI mice (area in mm^2^, sham: 1.11±0.06 in 8 mice, FPI: 1.20±0.09 in 6 mice, p=0.38 by Student’s t-test).

### Statistical Analysis

Statistical analyses were conducted in GraphPad Prism 9. The Shapiro-Wilk test was used to test for normality, and appropriate parametric or non-parametric statistical analyses were conducted. F-test to compare variances was used to assess whether both sample groups undergoing statistical comparisons have equal standard deviations. All statistical comparisons were conducted at an alpha level of α = 0.05. Two-way repeated measures ANOVA (TW-RM ANOVA), paired and unpaired Student’s or Welch’s t-test, Mann-Whitney U test and Kolmogorov-Smirnov (K-S) test were used where appropriate.

## RESULTS

### Dentate gyrus excitability is increased one week after concussive brain injury in mice

Previous studies have identified an early increase in dentate network excitability after concussive brain injury in rat (Santhakumar et al., 2001; Neuberger et al., 2017a; Korgaonkar et al., 2020b) and mice (Witgen et al., 2005; Smith et al., 2012; Folweiler et al., 2018; Folweiler et al., 2020). To determine if afferent input drive to GCs is enhanced after FPI in mice, we recorded granule cell current responses to stimulation of the perforant path fibers using an electrode placed outside the fissure in mice one week after FPI or sham injury. As illustrated by the IR-DIC images (Fig. 1A) and the lack of change in average cross-sectional area of the hilus (see methods), the moderate FPI used in this study did not result in gross anatomical distortions of the dentate gyrus. Recordings were obtained from a holding potential of −70 mV to isolate glutamatergic currents in the absence of synaptic blockers. As reported following FPI in rat (Korgaonkar et al., 2020b), perforant path-evoked excitatory post-synaptic current (eEPSC) amplitudes in response to increasing current injection, was significantly enhanced in mice one week after FPI compared to sham injured controls (Figure 1B-C, n = 14 cells from 5 sham mice and 8 cells from 4 FPI mice, F(1, 176)=31.25, p<0.0001 for effect of injury by TW-RM ANOVA). In light of the extensive changes in dentate inhibition after FPI (Gupta et al., 2012; Gupta et al., 2022), we isolated eEPSCs in the presence of saturating concentration of the ionotropic GABA_A_R antagonist gabazine (10µM). Granule cell eEPSC amplitude recorded in gabazine was also increased after FPI (Figure 1D-E, n = 11 cells/4 sham mice and 10 cells/4 FPI mice, F(1, 19)=15.49, p<0.0009 for effect of treatment by TW-RM ANOVA). Note that the prolonged eEPSC depolarization is consistent with polysynaptic activation previously reported after FPI in rat (Santhakumar et al., 2000). Analysis of the resting membrane potential (in mV, sham: − 65.63±2.95 in 12 cells, FPI: −64.7±1.92 in 10 cell, p=0.9 by t-test) and access resistance (in MΩ, sham: 14.1±1.4 in 12 cells, FPI: 15.3±1.5 in 10 cell, p=0.56 by t-test) failed to reveal systematic differences between groups. To assess whether FPI results in sustained increase in excitability one week after injury, we examined dentate sections for expression of ΔFosB, a uniquely stable activity-dependent immediate early gene product shown to be persistently enhanced in epilepsy (You et al., 2017). Immunohistochemical staining of sections from a cohort of sham and FPI mice sacrificed one week after FPI identified an increase in the number of ΔFosB expressing putative GCs in the granule cell layer compared to that in sham mice (Fig 1 F-G, number of ΔFosB + GCs: sham: 24.11±5.98, n = 8 mice FPI: 52.51±15.25, n = 7 mice, p=0.02 by Mann-Whitney U test), which is consistent with sustained enhancement of dentate excitability. There was an increase in variance in the number of ΔFosB expressing GCs after brain injury (F_6,7_ = 5.6, p=0.04) indicative of individual differences in dentate hyperexcitability. Together, these data establish that the mouse model of moderate FPI used in our study results in both an increase in excitatory drive to GCs and an overall sustained increase in dentate excitability one week after injury.

**Figure 1.**
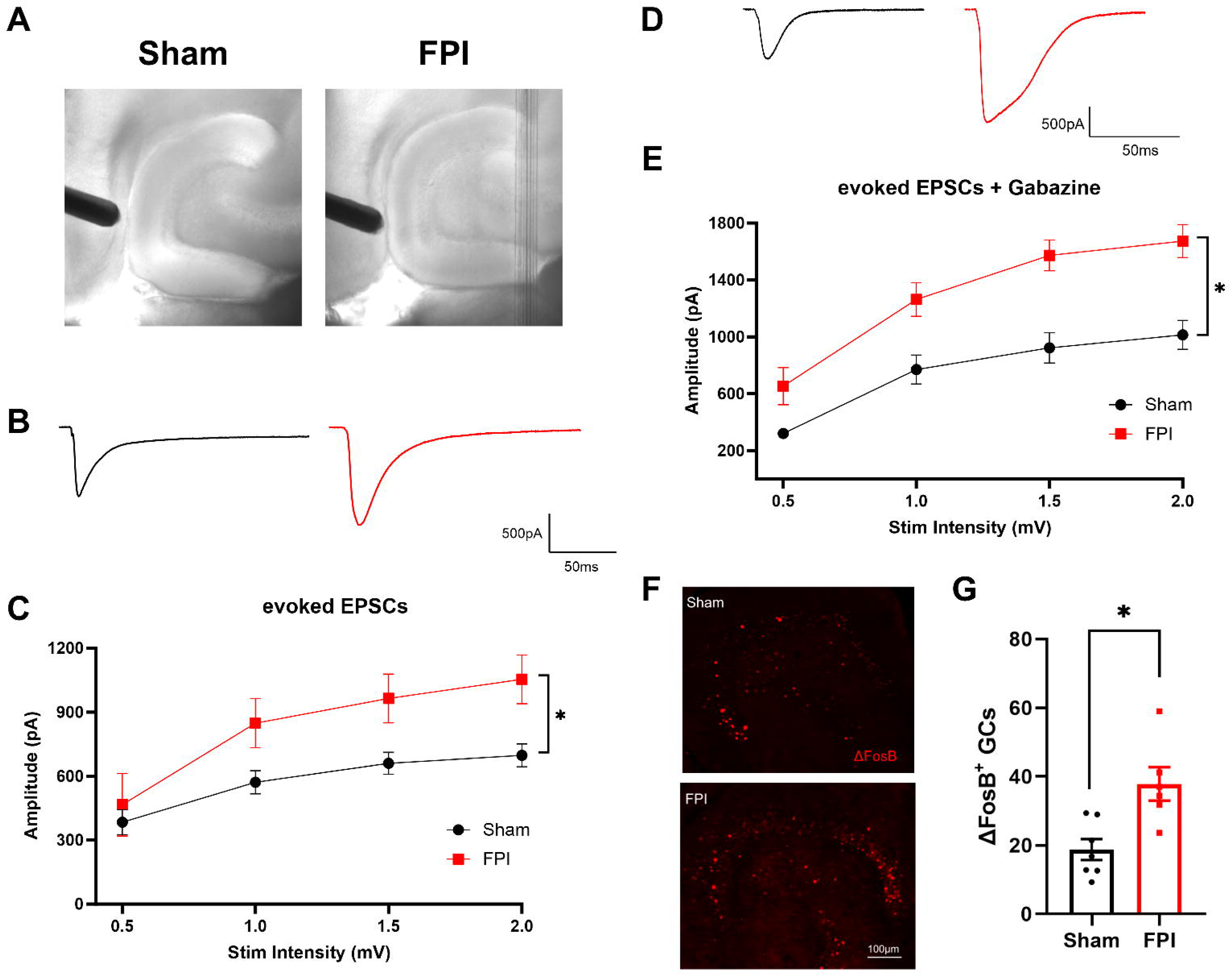
Enhanced dentate excitability one week after brain injury. (A) Representative IR-DIC images illustrating stimulating electrode positioning in slices from sham and FPI mice. (B) Representative eEPSC traces from dentate granule cells in sham (black) and FPI (red). (C) Plot of eEPSC peak amplitude in granule cells recorded in responses to increasing stimulus intensities. (D) Representative traces of pharmacologically isolated eEPSCs from dentate granule cells in sham (black) and FPI (red) recorded in 10 µM gabazine. (E) Summary plot of granule cell eEPSC peak amplitude in responses to increasing stimulus intensities recorded in 10 µM gabazine. (E). Representative images of staining for ΔFosB in sections (20 µm) from mice one week after sham and FPI. Scale bar=100µm. (F) Histogram showing average number of ΔFosB-positive cells in GCL of sham and FPI mice.

### Inhibition is critical for granule cell decorrelation of temporal spike train patterns

The characteristic low GC excitability and robust inhibition are critical for maintaining sparse GC firing and pattern separation. To determine if temporal input decorrelation by GCs is impacted by FPI, we implemented an ex-vivo temporal pattern separation paradigm (Madar et al., 2019a; Madar et al., 2021) to assess the ability of GCs, within the local dentate microcircuit, to disambiguate temporally patterned inputs. GC firing was recorded in response to stimulation of perforant path fibers in the outer molecular layer, using sets of stimulus patterns with known degrees of similarity, to quantify similarity between output spikes. For initial validation, four sets of Poisson input stimulus patterns with 10 Hz average frequency and input correlations of R25, R50, R75, or R95 were used to elicit GC responses (Fig 2A). Similarity indices of the GC “output” spike trains (R_out_) were computed as the pairwise Pearson’s correlation coefficient on output spike train data binned at 10, 20 and 100 ms to assess the effect of bin-width on R_out_ (Fig 2B). Consistent with Madar et al. (2019a), GC output similarity was consistently lower than the corresponding pairwise input similarity (Fig 2B, R_in_ vs. R_out_, n =4 cells / 3 mice). Since the reduction in output similarity was most robust when spike data were binned over 10 ms (Fig. 2B), a bin width of 10 ms was adopted in subsequent experiments. We then examined the potential contribution of the local inhibitory circuit to GC temporal pattern separation by partially decreasing GABAergic inhibition. To compare similarity indices before and after partial inhibitory block within the same cell, we restricted the inputs to a set of 5 stimuli with R75, reflecting a 75% similarity of across the inputs. GC responses were recorded first in aCSF and then in a non-saturating concentration of gabazine (100 nM). Under both recording conditions, GC output correlation remained consistently lower than the input correlation of 0.75 suggesting contributions from cell intrinsic and/or afferent synaptic characteristics in mediating GC temporal pattern separation. However, pairwise comparison of R_out_ within the same cell and pattern in aCSF versus gabazine revealed a significant increase in GC output correlation (Fig.2 C-E, Mean pairwise R_out_, aCSF: 0.20±0.02, Gabazine: 0.30±0.01, n = 4 cells /3 mice; p < 0.0001 by Paired t-test, Cohen’s D =1.7). These results validate the role for inhibition in the ability of GCs to decorrelate temporally patterned inputs as reported previously (Madar et al., 2019a).

**Figure 2.**
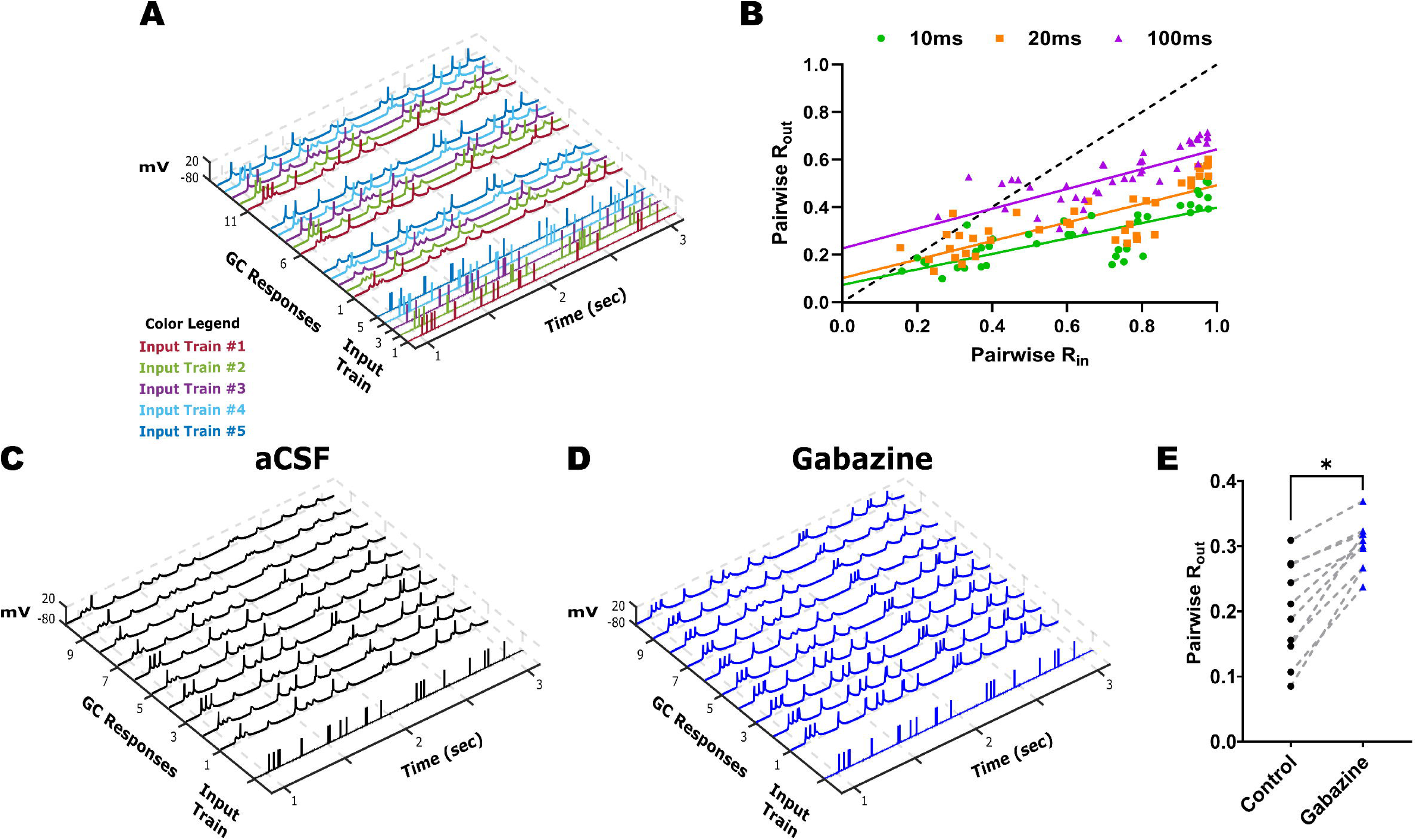
Robust temporal pattern separation by dentate granule cells. (A) Representative input spike trains from the R75 input set are illustrated with each pattern (input trains #1-5) in a distinct color. The GC firing evoked by three (of 10) repetitions of each input set is illustrated with output trace color matched to corresponding input spike train. (B) R_out_ vs R_in_ plots calculated with bin widths for correlation set to 10, 20, and 100 ms. Responses were elicited by input sets with similarity indices ranging from R25 to R95. (C-D) Example traces of GC firing responses elicited during 10 repetitions of the same representative (R75, pattern #4) input spike train recorded in aCSF (C) and in 100 nM gabazine (D). (E) Summary of pairwise R comparisons in aCSF and 100 nM gabazine (n = 4 cells). * Indicates p < 0.05 by Paired T-test.

### Reliability of dentate granule cell spiking is not altered early after FPI

After validating the experimental brain injury model in mouse (Fig. 1) and establishing the temporal pattern separation paradigm in slice recordings (Fig. 2), we examined GCs in mice one week after sham or FPI for their ability to decorrelate input spike train sets with 25% and 95% similarity. To ensure that there were no systematic differences in GC recordings between sham and FPI, we confirmed that the access resistance was not different between recordings from the two experimental groups (in MΩ, sham: 13.98±1.38, FPI: 15.57±1.97, p=0.54 by Student’s t-test). Additionally, examination of the GC intrinsic parameters failed to reveal differences in resting membrane potential (Fig. 3A-C, in mV, sham: −72.18±1.14, FPI: −75.86±2.06, p=0.16 by Student’s t-test) and input resistance (in MΩ, sham: 102.3±8.61, FPI: 121.1±13.34, p=0.28 by Student’s t-test, based on 11 cells/3 sham mice and 14 cells/4 FPI mice. In GCs that reached threshold with a +160 pA current injection, firing frequency was unchanged after FPI (in Hz, sham: 6.67±2.46 in 7 cells, FPI: 6.74±1.54 in 9 cells, p=0.98 by unpaired Student’s t-test). Similarly, action potential threshold measured at rheobase current injection was not different between groups (Fig. 3B, C, in mV, sham: −31.49±2.13, FPI: −32.15±1.78, p=0.81 by unpaired Student’s t-test). Given the neuronal loss and circuit alterations after FPI (Bonislawski et al., 2007; Neuberger et al., 2017a; Folweiler et al., 2020; Gupta et al., 2020), we examined whether the fidelity of GC response to a given input train is fundamentally altered by injury. Under optimal conditions, if a local dentate network receives identical inputs, a given GC within the receiving network should maintain indistinguishable output responses. To determine if injury alters the ability of the GC network to reliably respond to a given input train, we calculated spike train reliability (R_w_) as the average R_out_ of a given GC in response to identical input pattern using the R95 input set (Fig 4A, B). Note that because stimulus intensity was set to achieve a 50% spike success in each cell, the enhanced eEPSC (Fig. 1) should not impact evaluation of spike train reliability. Indeed, the average firing rate elicited by the input spike trains was not different between GCs from sham and FPI (Fig. 4C, average firing frequency in Hz, sham: 3.55±0.38, FPI: 2.99±0.19, p=0.17 by unpaired Student’s t-test). Interestingly, the variance of the firing frequency trended towards a reduction after FPI (F_10,13_= 3.10, p = 0.06, F-test for variance) suggesting a potential decrease in a source of input decorrelation after FPI. Despite the dentate circuit changes and enhanced ΔFosB labeling in GCs after FPI, spike train reliability was not different between the GCs from sham and FPI mice (Fig. 4C, average spike train reliability: sham: 0.47±0.05, FPI: 0.51±0.03, p=0.44 by unpaired Student’s t-test; variance (F_10,13_= 1.92, p = 0.27 by F-test for variance).

**Figure 3.**
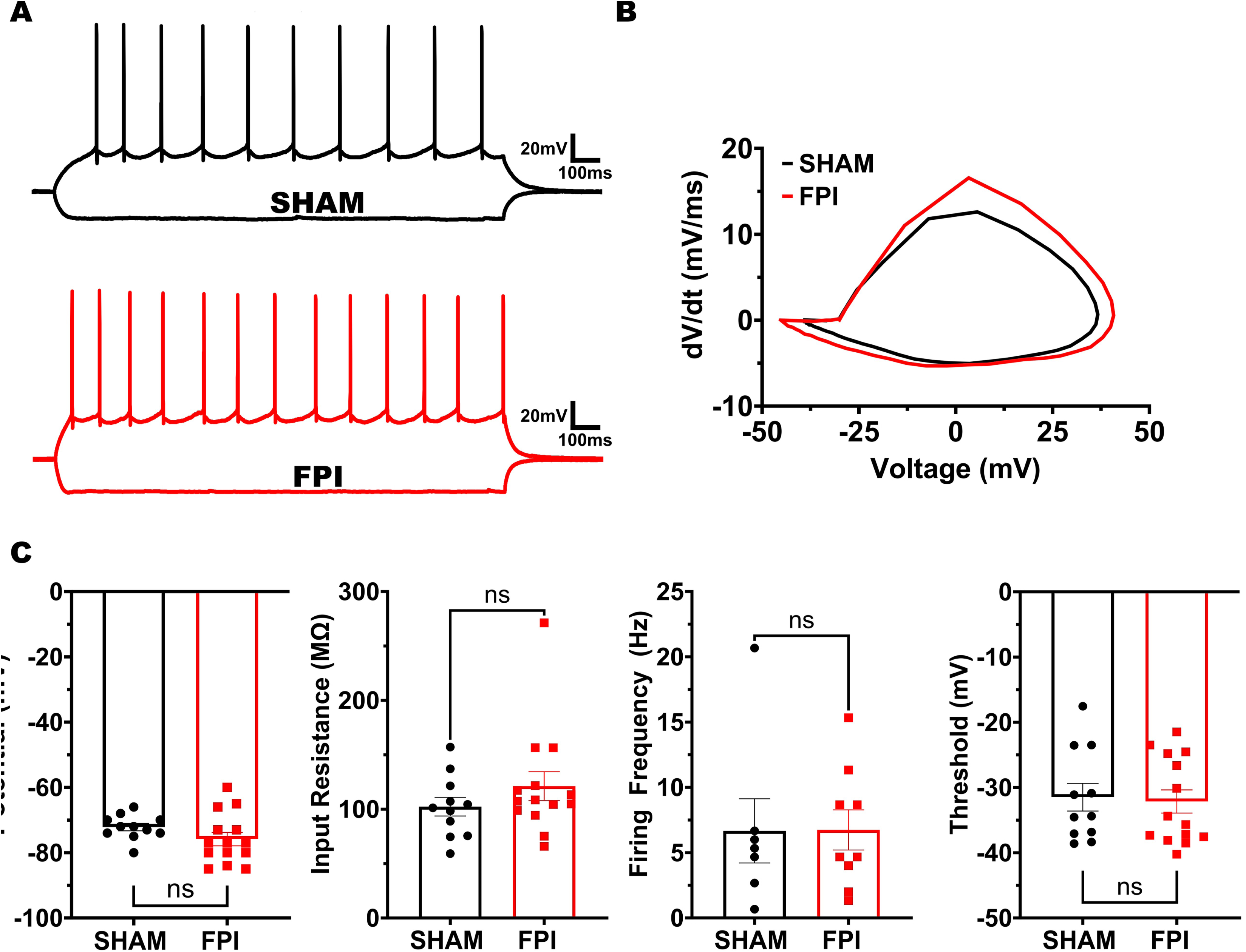
Granule cell intrinsic physiology is not altered FPI. (A) Representative granule cell membrane voltage traces in response to −200 pA and +160 pA current injections. (B) Representative action potential phase plots obtained from the first action potential in granule cells sham (black) and FPI (red) mice. (C) Summary plots of resting membrane potential, input resistance, firing frequency at +160pA i-inj and action potential threshold. Note that only cells that reached threshold at +160pA were included in analysis of firing frequency.

**Figure 4.**
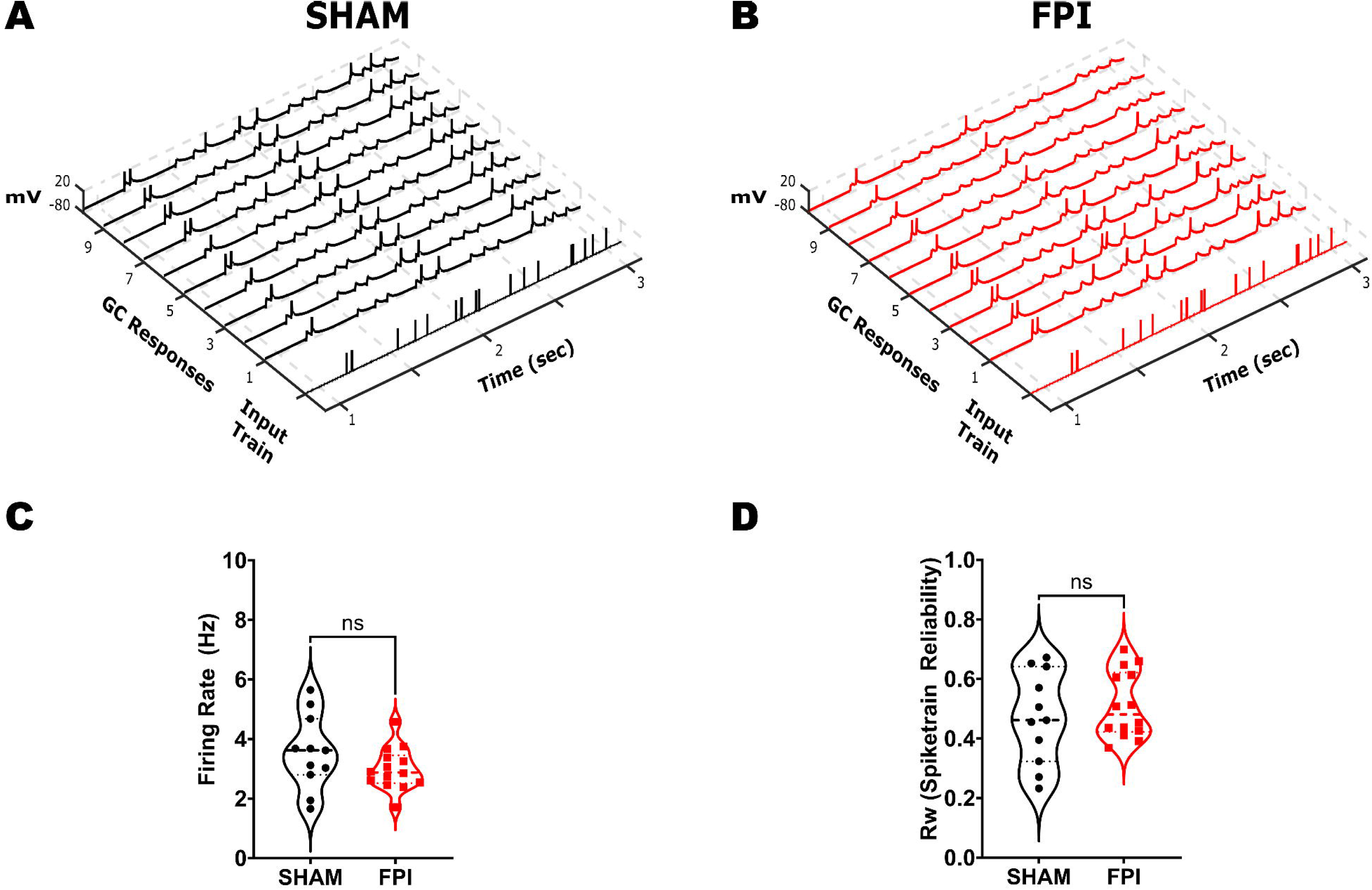
Assessment of GC firing fidelity after FPI. (A-B) Example of a single input spike train at R95 and the corresponding membrane voltage traces recorded in GC during 10 repetitions of the same stimulus in sham (A) and FPI (B) mice. (C-D) Summary plot of GC firing frequency (C) and reliability index (D) of GC firing during R95 inputs in sham (n = 11 cells from 3 mice) and FPI (n = 14 cells from 3 mice).

### Granule cell decorrelation of temporal patterned inputs is reduced after FPI

To compare temporal pattern separation between GCs from sham and FPI mice, GCs were stimulated using input stimulus train sets with 25% and 95% similarity (Fig. 5A-B). Plots of averaged pairwise R_out_ values against R_in_, for each input set showed that the GCs from both sham and FPI mice were still able to decorrelate inputs (R_out_ < R_in_) (Fig. 5C). Note that the input trains each average R_in_ generate 10 distinct pairwise R_in_ values, reflecting correlation between 2 input trains, with corresponding 10 pairwise R_out_ values for each cell (Supplementary Fig 1C, D). Comparison of the R_out_ values obtained in response to the 20 R_in_ values (10 each at R25 and R95) in GCs from sham and FPI mice revealed a significant effect of injury (F(1,18) = 6.57, p<0.02 by TW-RM ANOVA, based on 9 Sham cell from 3 mice and 11 FPI cells from 4 mice each where all input trains were recorded). However, the R_out_ within each cell averaged across the 10 R_in_ values at R25 (Sham: 0.13±0.01 in 9 cells/3 mice, FPI: 0.12±0.01 in 11 cells/4 mice, p=0.36) and R95 (Sham: 0.47±0.04 in 11 cells/3 mice, FPI: 0.49±0.01 in 14 cells/4 mice, p=0.49) showed no apparent difference (Supplementary Fig 1A, 1B, note that a few cells in which only R95 stimulus trains were recorded were included in this analysis). We reasoned that this was because averaging across multiple R_in_ values fails to capture the differences between R_in_ values and corresponding R_out_ (Supplementary Fig 1C, D). We generated cumulative probability distributions of the R_out_ data corresponding to each average R_in_ value to better assess changes in decorrelation. The cumulative probability distribution of all R_out_ values for R_in_ values centered around R25 was not different between GCs from sham and FPI (Fig. 5D, p=0.8 by K-S test). However, the distribution of all R_out_ values in response to inputs centered around R95 was significantly different between groups (Fig. 5E, p=0.0005 by K-S test) with a preferential reduction in the ability of GCs to support greater decorrelation after FPI. These findings are consistent with a subtle reduction in the ability of the dentate circuit to decorrelate highly similar inputs after injury. These data show that although GCs are continuing to disambiguate similar spike train inputs early after injury, their critical ability to do so when inputs are highly similar is compromised after injury.

**Figure 5.**
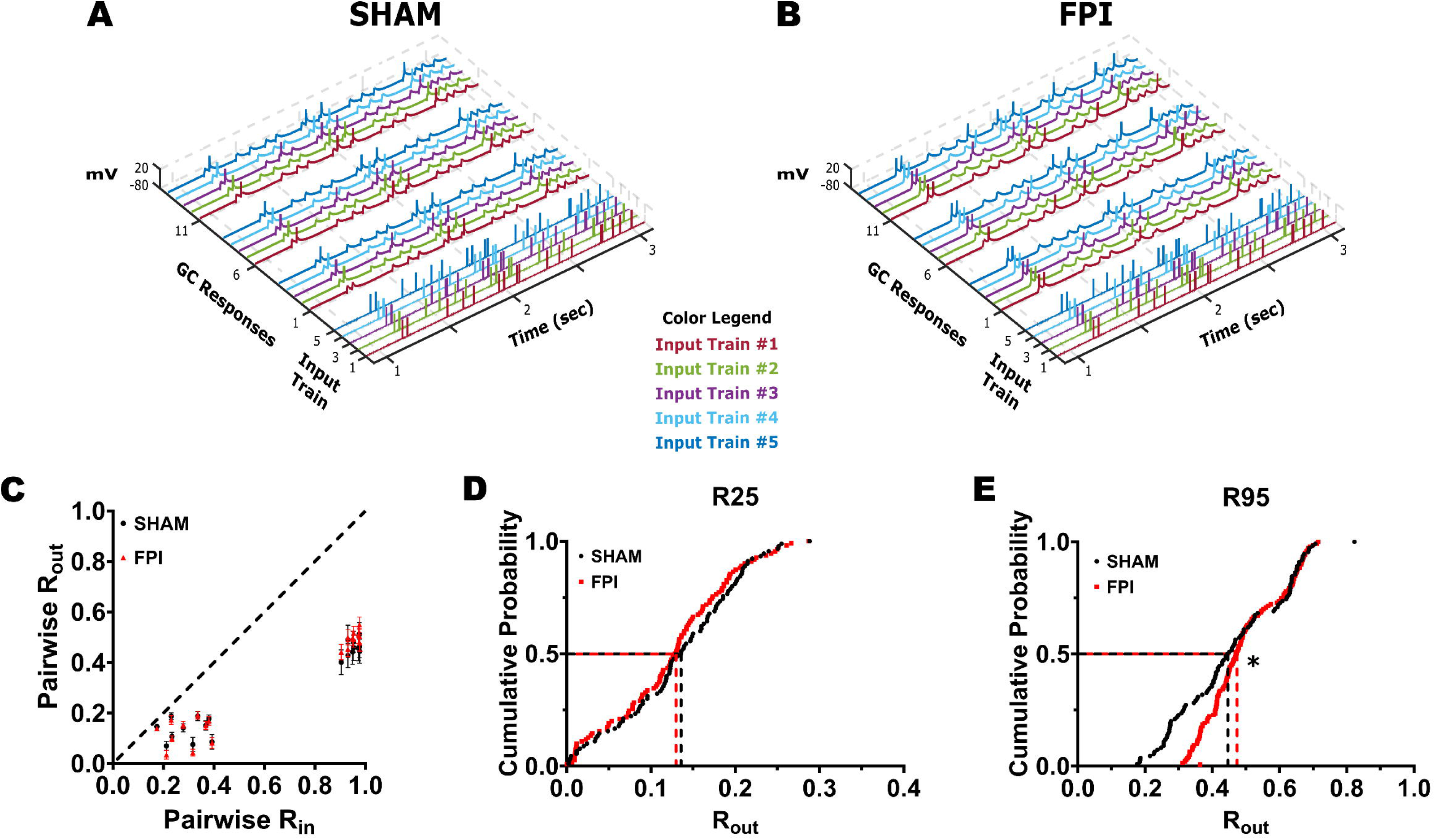
GC temporal pattern decorrelation is maintained after brain injury. (A-B) Representative input spike trains from the R95 input set are illustrated with each pattern (input trains #1-5) in a distinct color. GC firing evoked by three repetitions of each input set in sham (A) and FPI (B). Each membrane voltage trace is color matched to corresponding input spike train. (C) Summary plot of R_out_ vs R_in_ recorded during presentation of stimulus sets with R25 and R95 similarity indices in GCs from sham (black) and FPI (red) mice. (D-E) Cumulative probability distributions of pairwise R_out_ in GCs from sham and FPI mice during presentation of input sets with average R_in_=R25 (D, recordings from 9 cells in 3 sham mice and 11 cells in 3 FPI mice) and R95 (E, 11 cells in 3 sham mice and 14 cells in 3 FPI mice) * Indicates p < 0.05 by K-S test.

### Behavioral spatial pattern separation is degraded early after FPI

To determine whether the reduction in the ability of the dentate circuit to decorrelate input patterns after injury was accompanied by deficits in behavioral spatial discrimination, we adopted a novel object location task, which relies on dentate function and tests for episodic rather than sequential memory (Morales et al., 2021). Mice one week after FPI or sham surgery were assessed in the novel object location task following appropriate habituation (Fig. 6A). The mice were exposed to three identical objects in the sample phase followed, 30 minutes later, by a choice phase in which one object was removed and one of the remaining objects was moved to a novel location (Fig. 6A-B). Compared to sham mice, which showed significantly greater interaction with the object in the novel location (p=0.0002 by one sample t-test), FPI mice failed to show preference for the object in the novel location (p=0.67 by one sample t-test). Consistently, the discrimination ratio, defined as the difference between number of interactions with the object in the novel location and the familiar location divided by the total interaction with both objects, was lower in FPI mice (Fig. 6C, discrimination ratio: sham: 0.28±0.04, FPI: 0.04±0.08, n = 8 mice/group, p=0.02 by unpaired Student’s t-test). However, the total exploration assessed as the number of contacts with objects in the 5-minute exploration period was not different between groups (Fig. 6D, number of contacts: sham: 39.25±5.58, FPI: 41.13±6.6, n = 8 mice each, p=0.83 by unpaired Student’s t-test) indicating no difference in exploration between sham and FPI mice.

**Figure 6.**
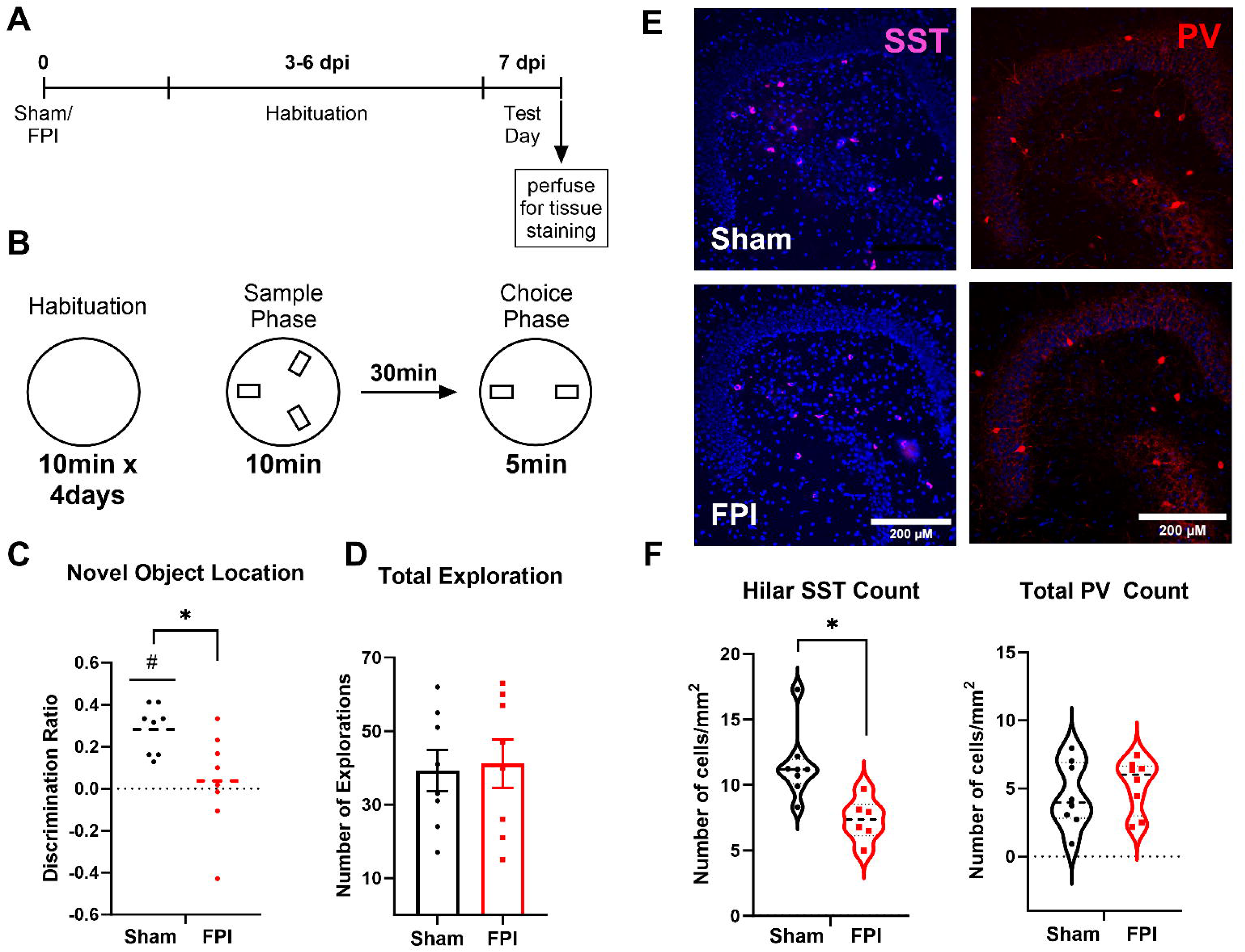
Spatial recognition memory is impaired following FPI. (A) Timeline of behavioral task and tissue processing for histological staining. (B) Schematic representation of the novel object location task performed one week post injury showing 4-day habituation in open chamber and test day timeline and location of objects in sample and choice phases. (C-D) Summary plot of discrimination ratio (C) and total duration of exploration (D) in sham (n = 8) and FPI (n=8) mice. (E) Representative images of dentate sections immunostained for somatostatin illustrates reduction in hilar somatostatin neurons one week after FPI, scale bar=100µm. (F) Summary histogram of somatostatin cell counts in sections from the dentate hilus in sham and FPI mice. (G) Representative images of dentate sections from sham and FPI mice immunostained for parvalbumin. (H) Summary histogram of PV cell counts/sections quantified in all dentate layers in sham and FPI mice.

Finally, because, previous studies have identified a role for dentate somatostatin interneurons in performance of the novel object location task (Morales et al., 2021) and somatostatin neurons are vulnerable to brain injury (Lowenstein et al., 1992; Santhakumar et al., 2000; Frankowski et al., 2019), we examined whether there was a loss of somatostatin neurons in the FPI mice that performed the behavioral task. Staining for somatostatin revealed a significant reduction of labeled neurons in the dentate hilus one week after FPI (Fig 6E-F, cells/mm^2^, sham: 11.50±0.92, FPI: 7.337±0.66, based on within animal averages of level matched sections from 8 sham and 6 FPI mice, p=0.005 by unpaired Student’s t-test). In contrast immunostaining for PV failed to reveal changes in PV neuron density after FPI (Fig 6E-F, cells/section, sham: 4.52±0.86, FPI: 5.22±0.70, based on within animal averages of level matched sections from 8 sham and 8 FPI mice, p>0.05 by unpaired Student’s t-test). These data suggest that injury-induced selective loss of dentate somatostatin interneurons could contribute to the observed impairment in location discrimination in mice early after brain injury.

## Discussion

Changes in episodic memory are a major neurocognitive consequence of traumatic brain injury with early and persistent deficits observed in patients and in animal models (Hamm et al., 1996; McHugh et al., 2006; Tsirka et al., 2010; Kumar et al., 2013; Smith et al., 2015; Korgaonkar et al., 2020a; Alosco et al., 2023). Although the dentate gyrus is known to play an essential role in encoding and disambiguating spatial memories, the underlying circuit mechanisms are not fully understood (Rolls and Kesner, 2006; Morris et al., 2012; Kesner, 2013; Cayco-Gajic and Silver, 2019). Here we implemented parallel circuit and behavioral assays of pattern separation in mice one week after concussive brain injury. Interestingly, we find that the reliability of GC firing is not altered after brain injury. Moreover, the GC output correlation was consistently lower than input spike train correlation in cells from both control and injured mice, indicating that temporal pattern separation at the cellular level continued to occur after brain injury. However, the critical ability of dentate GCs to decorrelate highly similar input patterns was reduced after brain injury. These results are analogous to the effect of partial GABA_A_ receptor antagonism where GC output correlation is higher than in aCSF, yet, remains lower than input correlation indicating deficits in decorrelation of inputs by GCs (Fig. 2 and Madar et al., 2019a). Despite the limited decrease in GC cellular temporal pattern decorrelation after injury, behavioral discrimination of spatial location in an episodic memory test was severely compromised. It is possible that the posttraumatic increase in the number of GCs showing sustained increase in excitability, identified by ΔFosB labeling, degrades sparse coding in the dentate and contributes to decline in spatial location discrimination. Furthermore, we demonstrate a decrease in dentate hilar somatostatin neurons in brain injured mice that showed deficits in spatial discrimination. Since somatostatin neurons have been shown to contribute to the novel object location task (Morales et al., 2021) used in our study, posttraumatic somatostatin neuron loss could contribute to deficits in behavioral spatial discrimination. Together these findings identify early changes in dentate inhibition and an increase in GC activity that could compromise the ability of the dentate to effectively decorrelate spike trains and engage distinct GC ensembles, likely contributing to deficits in encoding episodic memories.

### Posttraumatic changes in dentate circuit impact temporal decorrelation of input patterns

There is considerable evidence for the dentate gyrus playing a critical role in distinguishing context and spatial locations (Leutgeb et al., 2007; McHugh et al., 2007; Kesner, 2013; Senzai and Buzsaki, 2017). Because events occur in space and time, patterned neuronal firing can be essential for encoding spatial and temporal information (Kobayashi and Poo, 2004; Buzsaki, 2010; Tort et al., 2011; Eichenbaum, 2013). Experimental and computational studies examining dentate gyrus processing of temporally patterned inputs have identified that individual GCs can effectively orthogonalize temporal patterns in input spike trains. The ability of GCs to decorrelate temporal patterns likely depends on a combination of their intrinsic properties, local circuit inhibition, probabilistic synaptic release and spike-timing history resulting from short-term synaptic dynamics (Yim et al., 2015; Madar et al., 2019b, a). By adopting a temporal pattern separation paradigm, we confirmed that GCs effectively decorrelate input patterns, particularly those with high similarity (Fig. 2). Additionally, we show that even low dose gabazine (100 nM), resulting in partial block of inhibition, contributed to input decorrelation (Fig. 2) demonstrating a role for inhibition in decorrelation of input patterns. It is possible that, in addition to inhibition, perforant path synaptic dynamics and GC intrinsic physiology also contribute to temporal pattern separation in GCs.

In mice one week after concussive brain injury, we demonstrated an increase in perforant path evoked excitatory drive (Fig 1) which is consistent with our prior studies identifying an immune receptor mediated increase in AMPA currents one week after FPI in rat (Li et al., 2015; Korgaonkar et al., 2020b). These results and the lack of changes in GC intrinsic physiology (Fig. 3) and loss of hilar somatostatin neurons early after FPI are consistent with earlier studies in rats (Lowenstein et al., 1992; Santhakumar et al., 2000; Korgaonkar et al., 2020b). Moreover, labeling for ΔFosB, which indicates sustained increases in excitability (You et al., 2017), is increased one week after FPI. Therefore, it may seem surprising that the reliability and average firing frequency of GC responses to temporally patterned inputs remains unchanged after brain injury (Fig 4). Indeed, spike train reliability, a measure of the output correlation in response to identical inputs, is consistently lower than unity due to contributions from network noise and probabilistic transmitter release. A possible reason for maintenance of GC reliability despite the posttraumatic increase in afferent drive (Fig. 1) could be because, in the absence of post-injury changes in GC intrinsic physiology, setting the input stimulus intensity to elicit response in 50% of trials limited the effect of the increased excitatory input. Additionally, despite the loss of interneuron populations (Santhakumar et al., 2000; Folweiler et al., 2020), dentate inhibitory circuits undergo reorganization including increase in granule cell tonic GABA currents (Gupta et al., 2012; Gupta et al., 2022) which could help maintain GC activity levels. Moreover, GCs continue to effectively decorrelate input spike patterns as demonstrated by the consistent observation that output correlation remains lower than the input correlation even after brain injury (Fig 5C). Our finding that GCs continue to decorrelate input patterns early after brain injury is also consistent with recent findings in computational and experimental models of temporal lobe epilepsy (Yim et al., 2015; Madar et al., 2021). However, our results reveal a significant decrease in temporal pattern separation in response to input trains with high degrees of similarity, which suggests that the network level changes including a partial loss of hilar somatostatin neurons do impact decorrelation of input information. Because suppressing dentate inhibition with saturating concentrations of gabazine leads to burst firing in GCs (not shown), complete disinhibition is unlikely to yield meaningful analysis of decorrelation. Future studies adopting opto- or chemogenetic suppression of specific neuronal subtypes could provide additional insights into the contribution of specific interneuron subtypes to GC input decorrelation. Although injury did not decrease GC spike reliability, the variance in firing frequency in response to identical input trains trended to decrease after injury. It is interesting to speculate that the mechanisms contributing to reduced variance in GC firing after injury could also reduce GC decorrelation of highly similar inputs.

### Distinguishing between cellular decorrelation of spike timing and behavioral discrimination of temporally ordered memories

It must be noted that the temporal pattern separation paradigm used in this study focused on timing of spike trains and does not directly correspond to the encoding of temporal sequences of events. Indeed, it is unclear whether the dentate gyrus is involved in distinguishing the temporal order of sequential events occurring over a span of minutes to days (Gilbert et al., 2001). However, the dentate gyrus, and particularly the ongoing generation of adult-born neurons in the dentate, has been shown to play a role in stratification of memories over the time course of weeks (Clelland et al., 2009; Aimone et al., 2011; Nakashiba et al., 2012; Rangel et al., 2014; Miller and Sahay, 2019). Interestingly, adult neurogenesis is altered after brain injury with early increase at one week followed by a progressive decline at one month (Yu et al., 2008; Kernie and Parent, 2010; Shapiro, 2016; Neuberger et al., 2017a; Ngwenya and Danzer, 2018; Clark et al., 2020). Whether injury induced changes in neurogenesis alter stratification or timestamping of contemporary memories and/or compromise spatial and contextual pattern separation at later time points after injury remains to be examined. In this study we focused on the early time point, one week after injury, when the injury-induced neurons are believed to be immature and unlikely to contribute to local dentate feedback inhibition (Overstreet-Wadiche and Westbrook, 2006; Temprana et al., 2015; Miller and Sahay, 2019). Additionally, we recorded GCs in the middle and outer third of the granule cell layer so as to avoid immature adult-born neurons, typically located in the inner third of the granule cell layer (Save et al., 2018; Kerloch et al., 2019). Crucially, all GCs included in analysis had input resistance less than 300 MΩ consistent with mature GC phenotype. Thus, our results identify that, at the level of individual putative mature dentate GCs, spike reliability and ability to decorrelate spike train patterns persist despite the early circuit changes after brain injury. Still, considering that temporal correlations between dentate neurons at the sub-second time scale are important for spatial memory discrimination (van Dijk and Fenton, 2018), our slice physiology data suggest that the dentate’s ability to encode input timing is largely retained early after brain injury. Because the dentate gyrus is particularly important for disambiguating highly similar patterns, the posttraumatic reduction in GC ability to decorrelate highly similar inputs could contribute to deficits in pattern separation after brain injury.

### Mechanisms underlying altered behavioral pattern separation after brain injury

In contrast to GC temporal pattern separation, we find that the ability to encode and discriminate subtle differences in spatial location is profoundly compromised one week after brain injury (Fig. 5). Our results complement a previous study which showed that mice one week after mild to moderate concussive brain injury are impaired in a sequential location discrimination task using the radial arms maze (Correll et al., 2021). Unlike the radial arm maze, the novel object location adopted in our study examines episodic memory and does not rely on repetition of discrete trial procedures (Bekinschtein et al., 2013; Morales et al., 2021). At the medium difficulty level (60° relocation) adopted in our study, the task has been shown to rely on dentate processing and is compromised by selective suppression of dentate somatostatin neurons. Consistently, we find that the behavioral deficit in novel object location one week after brain injury is accompanied by selective loss of hilar somatostatin neurons suggesting that interneuron loss may drive deficits in spatial pattern separation. Our data showing lack of change in PV neurons one week after FPI, when quantified across all dentate subregions, is consistent with Folweiler et al. (2020). However previous studies have identified post-FPI loss of PV neurons restricted to the dentate hilus and altered inhibition after FPI (Santhakumar et al., 2000; Folweiler et al., 2020). Whereas loss of immature and highly excitable adult born neurons born prior to injury could contribute to deficits in encoding novel object location, studies using the cortical impact injury have reported no loss of GCs born prior to injury (Kang et al., 2022). Rather, these neurons support enhanced feedback inhibition 8-10 weeks after injury. Moreover, it should be noted that neurogenesis is increased rather than decreased one week after injury (Shapiro, 2016; Neuberger et al., 2017a; Clark et al., 2020; Correll et al., 2021) and adult born neurons support local circuit feedback inhibition of mature GCs 4-6 weeks into development (Temprana et al., 2015; Miller and Sahay, 2019). Taken together, it is unlikely that injury induced changes in dentate neurogenesis underlie deficits in spatial pattern separation observed one week after injury. Instead, post-injury reduction of feedback inhibition, due to loss of somatostatin neurons, and increases in basal activity of GCs, revealed by the enhanced ΔFosB staining, likely impair the ability of the dentate circuit to effectively decorrelate entorhinal inputs into discrete active GC ensembles and firing patterns.

### Reconciling the modest loss in cellular temporal decorrelation with profound impairment in behavioral spatial discrimination

The apparent divergence in effect of brain injury on decorrelation of spike trains at the cellular level and discrimination of spatial location at a behavioral level indicate that disambiguation of information likely involves activation of distinct neuronal ensembles by input patterns. This is consistent with findings in computational models (Myers and Scharfman, 2009, 2011). Experimentally, due to the time for genetically encoded reporters to express (Guenthner et al., 2013; Ramirez et al., 2013), it may be challenging to identify neurons activated by two distinct and recent spatial inputs. However, there is some evidence for sparse but shared activation of neurons by different spatial experiences (Tashiro et al., 2007). The basal increase in neuronal excitability, identified by increase in ΔFosB labeled GCs after brain injury, could corrupt this sparse and selective activity necessary for discrimination of spatial memories. A potential caveat is that, whereas the circuit experiments focused on the outer molecular layer, which receives the lateral entorhinal cortical fibers carrying contextual information, the spatial information examined in the behavioral spatial discrimination study is thought to be carried by inputs from the medial entorhinal cortex. Future studies examining decorrelation of medial perforant path inputs and behavioral analysis of context encoding could resolve whether the differences identified here are pathway specific. Analysis of spatiotemporal activity patterns of neuronal ensembles during behaviors or imaging circuit activity in vitro in response to spatiotemporal input patterns would help resolve how the dentate disambiguates input patterns.

In summary, our study identifies that early posttraumatic changes, including persistent increase in excitability and loss of somatostatin interneurons, reduce temporal pattern separation by dentate GCs and degrade the ability to discriminate between similar spatial locations. These changes likely contribute to working memory deficits after brain injury.

## Supporting information

Corrubia-Supplementary Materials

## Acknowledgements

We thank Dr. Krista Marrero for thoughtful comments and discussion.

## Author Contributions

L.C. A.H L.A.E and S.N. Investigation; Formal analysis; Data curation; Methodology; Figures; A.H. Software; L.C. A.H., S.N. M.W.S., M.V.J., L.A.E and V.S. interpreted results of experiments; A. H., L.C. and S.N. prepared figures; A.H., L.C., S.N., and V.S. Conceptualization V.S. Funding acquisition; Project administration; Resources; Supervision; L.C. A.H., S.N. M.W.S., M.V.J., L.A.E and V.S. Writing - review & editing.

## Supplementary Figures

**Supplementary Figure 1. Decrease in GC decorrelation of highly similar temporal input patterns after brain injury.** (A-B) Summary plots of average R_out_ in each cell averaged across all R_in_ values centered around R25 (A) and R95 (B). (C-D) Pairwise Rout/Rin in GCs from sham and FPI mice for 10 distinct Rin values each centered around R=0.25 (C) and R=95 (D). Note that these are the same data as in Fig 5C presented at a different scale.

