## Supplementary material for "Early Deficits in Dentate Circuit and Behavioral Pattern Separation after Concussive Brain Injury": Corrubia-Supplementary Materials

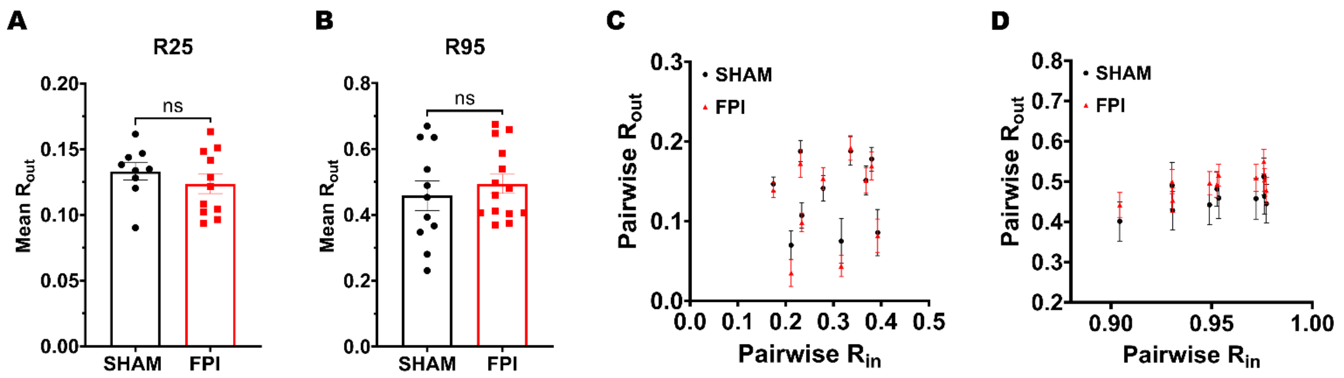


**Supplementary Figure 1. Decrease in GC decorrelation of highly similar temporal input patterns after brain injury.** (A-B) Summary plots of average Rout in each cell averaged across all Rin values centered around R25 (A) and R95 (B). (C-D) Pairwise Rout/Rin in GCs from sham and FPI mice for 10 distinct Rin values each centered around R=0.25 (C) and R=95 (D). Note that these are the same data as in Fig 5C presented at a different scale.
